# Cheating emergences in the arbuscular mycorrhizal mutualism: a network and phylogenetic analysis

**DOI:** 10.1101/500504

**Authors:** Benoît Perez-Lamarque, Marc-André Selosse, Maarja Öpik, Hélène Morlon, Florent Martos

## Abstract

- While mutualisms are widespread and essential in ecosystem functioning, the emergence of uncooperative cheaters threatens their stability, unless there are functional or evolutionary mechanisms limiting cheaters interactions.
- Here, we evaluated the constraints upon mycoheterotrophic (MH) cheating plants in the mutualistic interaction network of autotrophic (AT) plants and arbuscular mycorrhizal fungi. For this purpose, we assembled a world-scale network of >25,000 interactions in order to investigate *(i)* the specialization and *(ii)* the phylogenetic distribution of MH or AT plants and their respective fungal partners.
- We show that MH cheating repeatedly evolved in the vascular flora, suggesting low phylogenetic constraint for plants. However, MH cheaters are significantly more specialized than AT plants, and their fungi also appear more specialized and more closely related than fungi of AT plants, which suggest that cheaters are specifically isolated into modules by functional constraints
- This unprecedented comparison of MH vs. AT plants thus reveals that MH cheating is most likely constrained by the specialization of phylogenetically conserved cheating-susceptible fungi, which suggests mechanisms for avoidance of these fungi. Beyond the mycorrhizal symbiosis, our approach highlights an empirical multiple-partners mutualistic system illustrating that the overall persistence of mutualism can be linked to functional constraints upon cheating emergences.

## Introduction

Mutualistic interactions are ubiquitous in nature and largely contribute to generate and maintain biodiversity (Bronstein, 2015). Since benefits in mutualism often come at a cost for cooperators, some species called *cheaters* evolved an adaptive uncooperative strategy by retrieving benefits from an interaction without paying the associated cost (Sachs *et al.*, 2010). Although cheating compromises the evolutionary stability of mutualisms (Ferriere *et al.*, 2002), its evolutionary emergence is often limited by factors securing the persistence of mutualism (Bronstein *et al.*, 2003; Frederickson, 2013; Jones *et al.*, 2015). For instance, cooperators often favor the most rewarding partners (e.g. conditional investment; Roberts & Sherratt, 1998), stop interactions with cheaters (Pellmyr & Huth, 1994), or even sanction them (Kiers et al. 2003). Cheating emergence can thus be constrained through physiological or biochemical mechanisms of the interaction and its regulation, a type of functional mechanisms hereafter referred to as *functional constraints*. In addition, these functional constraints, and potentially others such as geographic constraints, can either be evolutionary conserved or not. If they are, there will be *phylogenetic constraints* on the emergence of cheaters, as some species will be evolutionary predisposed to cheating, or predisposed to be cheated upon, while others will not (Gómez *et al.* 2010).

The framework of bipartite interaction networks, combined with the phylogeny of partners, is useful for understanding the constraints limiting the emergence of cheaters in mutualisms. Analyses of bipartite networks have been extensively used to showcase the properties of mutualistic interactions (Bascompte *et al.*, 2003; Rezende *et al.*, 2007; Martos *et al.*, 2012), such as their level of specialization (number of partners), nestedness (do specialists establish asymmetric specialization with partners that are themselves generalists?), and modularity (existence of distinct sub-networks; Bascompte & Jordano 2014). These studies have shown that mutualistic networks are generally nested with specialists establishing asymmetric specialization with more generalist partners, contrary to antagonistic networks that tend to be modular, with partners establishing reciprocal specialization (Thebault & Fontaine, 2010). However, few analyses of bipartite networks include cheaters and analyze their specialization, nestedness and modularity (Fontaine *et al.*, 2011). Joffard *et al.* (2018) showed that specialization of orchids toward pollinators was higher in deceptive cheaters than in cooperative nectar-producing species. Considering the overall effect of cheaters on network structure, Genini *et al.* (2010) showed that a network dominated by cooperative pollinators was nested, whereas another network dominated by nectar thieving insects was more modular. If cheaters specialize and form modules, this suggests that functional constraints limit the set of species that they can exploit. Additionally, if cheaters emerged only once on a phylogeny (*versus* repetitively), and/or if ‘cheating-susceptible’ partners are phylogenetically related (Merckx *et al.*, 2012), this suggests that cheating involves some rare evolutionary innovations (Pellmyr *et al,.* 1996) and/or that cheating-susceptibility is limited to few clades, meaning that cheating is phylogenetically constrained.

Here we evaluate functional and phylogenetic constraints upon cheating in the arbuscular mycorrhizal mutualism between plant roots and soil Glomeromycotina fungi (Selosse & Rousset, 2011; Jacquemyn & Merckx, 2019). This 400 Myr-old symbiosis concerns ca. 80% of extant land plants and several hundred fungal taxa (Davison *et al.*, 2015; van der Heijden *et al.*, 2015). Arbuscular mycorrhizal fungi colonize plant roots and provide host plants with water and mineral nutrients, in return for carbohydrates. Although obligate for both partners, this symbiosis is generally diffuse and not very specific (van der Heijden *et al.*, 2015), since multiple fungi colonize most plants, while fungi are usually shared among surrounding plant species (Verbruggen *et al.* 2012). Thus, fungi interconnect plant individuals of different species, and allow resource movement between plants (Selosse *et al,.* 2006). This allowed the emergence of achlorophyllous cheating plants, called mycoheterotrophic (MH) plants, which obtain carbon from their mycorrhizal fungi that are themselves fed by surrounding autotrophic plants (AT) (Selosse & Rousset 2011; Merckx, 2013). Some of these plant species are entirely MH (*EMH*) over their lifecycle, while others are MH only at early stages before turning AT (initially MH, *IMH*), therefore shifting from cheater to potentially cooperative (Merckx, 2013). Unlike other cheating systems where cheating cost affects mostly direct partners (e.g. in plant pollination), MH cheaters involve not only a direct cost to their fungal partners but also a projected cost (i.e. a cost transmitted through the network) on the interconnected AT plants, whose photosynthesis supplies the carbon.

We evaluate functional constraints on MH by analyzing specialization, nestedness and modularity in the plant mycorrhiza interaction network. MH plants are thought to be specialists interacting with few fungal species (Leake, 1994; Merckx, 2013), but whether or not these species are unusually specialized compared to AT plants is still debated (Merckx *et al.*, 2012). MH plants could specialize on few fungal species if functional constraints limit the set of fungi they can exploit, and evolve more efficient nutritional stealing strategies (Blüthgen *et al.*, 2007). In terms of nestedness and modularity, arbuscular mycorrhizal networks are generally nested (Chagnon *et al.*, 2012), but how MH plants affect their structure has yet to be investigated. Because MH species indirectly rely on AT plants for carbon nutrition, interacting with generalist fungi could provide access to various carbon sources, and this would increase nestedness. Conversely, AT plants could sanction interactions with fungi associated with MH plants because of the projected cost; this would isolate cheating interactions within antagonistic modules and increase modularity. Establishment of extreme reciprocal specialization between one MH plant and one fungus seems unlikely though, since a AT carbon source is required.

With regards to phylogenetic constraints on MH, we already know that MH cheating strategies evolved multiple times (Merckx, 2013), generating monophyletic groups of MH plants, which suggests weak phylogenetic constraints on the emergence of cheating in plants. However, it seems that fungi interacting with independent MH plant families are phylogenetically close, suggesting that there are phylogenetic constraints on the fungal side (Merckx *et al.*, 2012). This needs to be confirmed in a larger phylogenetic context including the fungi of AT plants. Indeed, if a set of phylogenetically close fungi interact with all MH lineages, an important follow-up question is whether these fungi were derived independently from AT ancestors, or whether they were acquired by symbiont shift from other MH plants during new MH emergences.

We assess functional and phylogenetic constraints upon the emergence of MH cheaters in the arbuscular mycorrhizal mutualism using a combination of network and phylogenetic analyses. We analyze the largest available database on arbuscular mycorrhizal fungi and their host plants (Öpik *et al.*, 2010), which we updated for this analysis (see methods), and which includes both AT plants and most MH plant lineages.

## Materials and Methods

### Analytical framework

Figure 1 summarizes the main analyses we carried to evaluate functional and phylogenetic constraints upon the emergence of MH plants. We first assembled a bipartite network representing which plants interact with which fungi. Next, we analyzed functional constraints using tools from ecological networks (Fig. **1b**). Functional constraints on MH cheaters (or MH-associated fungi) should be reflected by the specialization of MH cheaters (or MH-associated fungi) towards few fungal (or plant) partners. Strong constraints on both sides should drive reciprocal specialization and isolation of MH cheaters and MH-associated fungi into modules, thus decreasing nestedness. Finally, we analyzed phylogenetic constraints by analyzing the phylogenetic distributions of MH cheaters and MH-associated fungi (Fig. **1a**).

**Figure 1:**
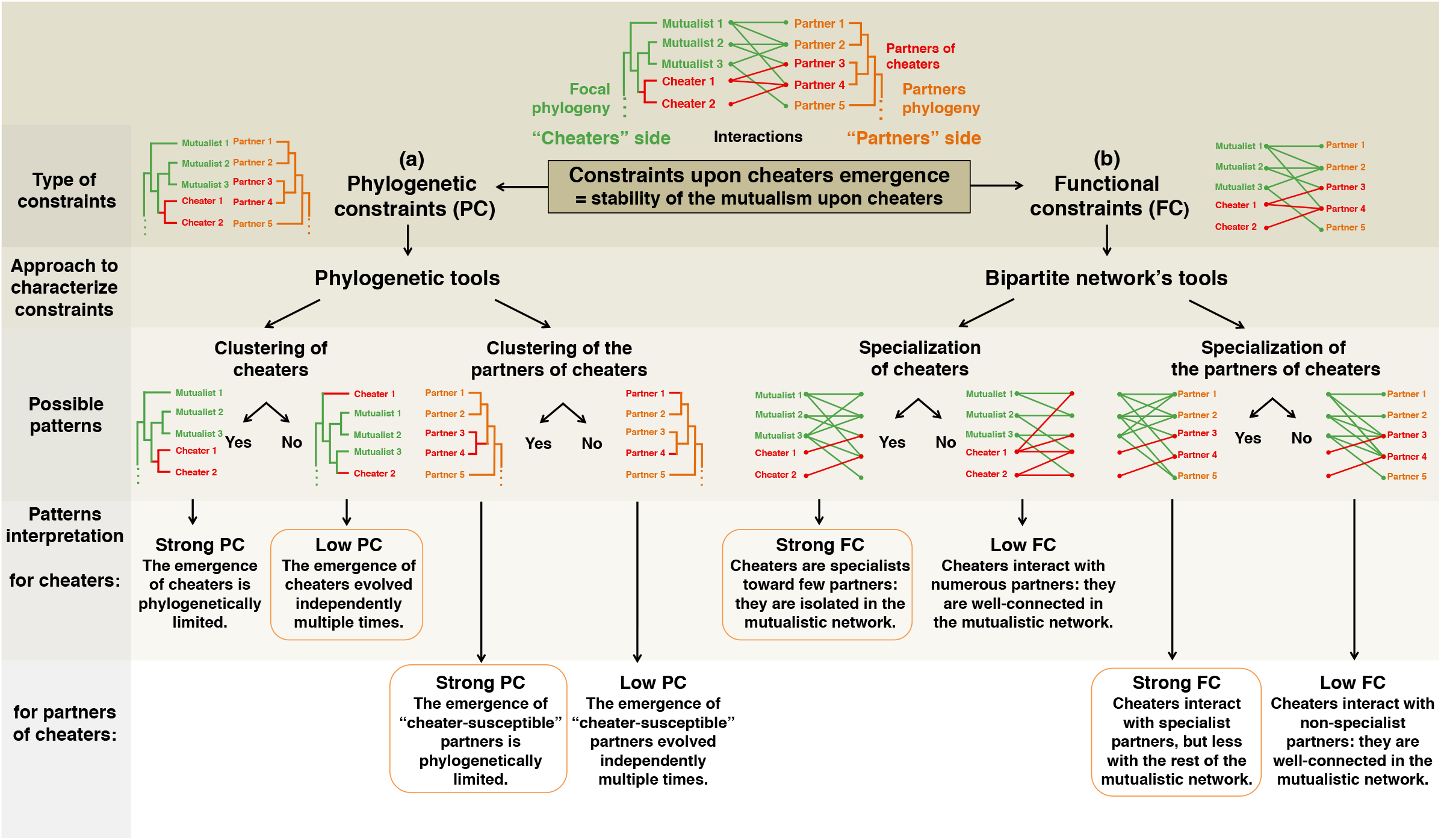
Framework used in this study to evaluate the constraints upon the emergence of mycoheterotrophic (MH) cheaters in the arbuscular mycorrhizal symbiosis. *(a)* Strong phylogenetic constraints (PC) should affect the phylogenetic distributions of MH cheaters and/or their fungal partners; whereas *(b)* functional constraints (FC) should affect the network structure i.e. level of specialization of MH cheaters and/or their partners. Therefore, by investigating specialization and phylogenetic clustering of MH cheaters and of their fungal partners, we evaluated functional and phylogenetic constraints. The patterns and interpretations from the present study are shown in the orange frames.

### MaarjAM database and interaction matrix

MaarjAM database is a web-based database (http://maarjam.botany.ut.ee; accessed in June 2019 after a very recent update) of publicly available sequences of Glomeromycotina fungi, with information on the host plants, geographical location and biomes for the recorded interactions (Öpik *et al.*, 2010, Davison *et al.*, 2015). We used an approach with a compiled network, where all locally described physical mycelial interactions between species are merged and studied at larger scales (as in Joffard *et al.*, 2018; Braga *et al.*, 2018). Although such a compiled network can be sensitive to several biases (see Discussion), it is a unique opportunity to study MH cheating emergence in a large evolutionary and ecological picture (e.g. Werner *et al.*, 2018). Among the 41,989 interactions between plants and Glomeromycotina, we filtered out the data from MaarjAM for the fungi to satisfy the following criteria (Supporting Information Table S1): *(i)* amplification of the 18S rRNA gene, *(ii)* fungus identified from plant roots (i.e. excluding soil samples and anthropogenic sites), *(iii)* interaction in a natural ecosystem (i.e. excluding anthropogenic or highly disturbed ecosystems), *(iv)* host plant identified at the species level, and *(v)* a virtual taxon (VT) assignation available in MaarjAM. The VTs are a classification (= species proxy) of arbuscular mycorrhizal fungi designed by applying a ≥97% sequence similarity threshold to the 18S rRNA sequences, and by running phylogenetic analysis to ensure VT monophyly (Öpik *et al.*, 2013, 2014). The filtered dataset yielded a binary interaction matrix of 490 plant species (hereafter ‘plants’), 351 VT (hereafter ‘fungi’), and 26,350 interactions, resulting from the compilations of 112 publications from worldwide ecosystems (Supporting Information Fig. **S6**; Supporting Information Table **S1**). In order to estimate the sampling fraction of Glomeromycotina fungi in our dataset, we performed rarefaction curves of the number of fungal VTs as a function of the sampling fraction (on the observed number of interactions or on the number of sampled plant species) and we estimated the total number of units using the ‘*specpool’* function (‘*vegan’* R-package, based on Chao index; Oksanen *et al.*, 2019).

### Phylogenetic reconstructions

We aligned consensus sequences of the 351 fungi with the software MUSCLE (Edgar, 2004), and ran a Bayesian analysis using BEAST2 to reconstruct the fungal phylogeny (Bouckaert *et al.* 2014, see Supplementary methods). We obtained the phylogenetic relationships between the 490 host plants by pruning the time-calibrated supertree from Zanne *et al.* (2013) using Phylomatic (http://phylodiversity.net/phylomatic/). We also used the Open Tree of Life website (http://opentreeoflife.org) and the ‘*rotl*’ package (Michonneau et al. 2016) of R (R Core Team, 2019) for grafting 41 plant taxa missing in the pruned supertree (as polytomies at the lowest taxonomy level possible, see Supplementary methods). We set tree root calibrations at 505 million years (Myr) for the fungi (Davison *et al.*, 2015) and 440 Myr for the plants (Zanne *et al.*, 2013).

### Nature of the interaction

We assigned to each plant its ‘nature of the interaction’ with fungi according to its carbon nutrition mode: autotroph (AT, n=434, 88.6%), entirely mycoheterotroph (EMH, n=41, 8.4%), or initially mycoheterotroph (IMH, n=15, 3.1%). According to an on-line database (http://mhp.myspecies.info/category/myco-heterotrophic-plants/) and individual publications (Boullard, 1979; Winther & Friedman, 2008; Field et al. 2015), EMH species belong to six monophyletic families [Burmanniaceae (25 spp.), Gentianaceae (6 spp.), Triuridaceae (4 spp.), Polygalaceae (4 spp.), Corsiaceae (1 sp.), and Petrosaviaceae (1 sp.)], and IMH species belong to three families [Ophioglossaceae (ferns; 5 spp.), Psilotaceae (ferns; 2 spp.), and Lycopodiaceae (clubmoss; 8 spp.)]. We assigned each fungus to three categories: ‘AT-associated’ if the fungus interacts with AT plants <u>only</u> (n=280, 79.8%), ‘EMH-associated’ if the fungus interacts with <u>at least</u> one EMH plant (n=54, 15.4%), ‘IMH-associated’ if the fungus interacts with <u>at least</u> one IMH plant (n=23, 6.6%), or ‘MH-associated’ if the fungus interacts with <u>at least</u> one EMH or IMH plant (n=71, 20.2%; Supporting Information Table **S2**). Only five MH-associated fungi are both EMH- and IMH-associated. Although our dataset unfortunately does not include all MH plants (it includes 18 publications), it covers a fair representative range of the mycoheterotrophs in the arbuscular mycorrhizal symbiosis (Jacquemyn & Merckx, 2019).

### Network nestedness and specialization of cheaters

In order to assess functional constraints, we tested the effect of MH cheating on network structure. First, we measured nestedness in: *(i)* the overall network (490 plants, 351 fungi, and 26,350 interactions), *(ii)* the network restricted to AT plants only (434, 344, and 26,087) and *(iii)* the network formed by EMH and IMH plants only (56, 71, and 263), using the function ‘*NODF2*’ in the R package ‘*bipartite’* (Dormann et al. 2008). Positive NODF values (nestedness metric based on overlap and decreasing fill; see *List of abbreviations* in Supplementary material) indicate nested networks. We tested the significance of nestedness values by using two types of null models (*N*= 100 for each type): both maintain the connectance (proportion of observed interactions) of the network, but the first model (*‘r2dtable’* from *‘stats’* R package - *null model 3*) also maintains the marginal sums (the sums of each row and each column) while the less stringent second model (*‘vaznull’* from ‘*bipartite’* R package - *null model 2*) produces slightly different marginal sums (interactions are randomized with species marginal sums as weights, and each species must have at least one interaction). We calculated the Z-score, which is the standard deviation between the observed value and the mean of the distribution of null-model networks (Z-scores greater than 1.96 validates a significant nestedness with an alpha risk of 5%).

Second, to further evaluate the specialization of MH cheaters, we computed several network indices for each plant. The degree (*k*) is the number of partners with which a given plant or fungus interacts in the bipartite network. The degree is high (resp. low) when the species is generalist (resp. specialist). The partner specialization (*Psp*) is the mean degree (*k*) averaged for all the fungal partners for a given plant species (Taudiere *et al.*, 2015): a high (resp. low) *Psp* characterizes a species interacting mainly with generalist (resp. specialist) partners. Simultaneously low *k* and *Psp* values feature a reciprocal specialization. Then, in order to detect specialization at clade scale toward partners, for any given clade of every node in the plant or fungus phylogenies, we calculated the partner fidelity (*Fx*) as the ratio of partners exclusively interacting with this particular clade divided by the total number of partners interacting with it. We consider the clade as ‘faithful’ and the corresponding set of partners as ‘clade-specific’ when *Fx*>0.5 (i.e., more than 50% exclusive partners). We tested whether *k* and *Psp* were statistically different among AT, EMH and IMH plants using non-parametric Kruskal-Wallis tests and pairwise Whitney U tests. We also replicated the statistical tests without the IMH Lycopodiaceae, because of their different network patterns (see Results). We used an analysis of covariance (ANCOVA) to test the effect of the nature of the interaction on partner fidelity *Fx* accounting for clade size, which corrects the bias of having high partner fidelity *Fx* in older clades including many plants.

To assess the significance of *k* and *Psp* values, we built null-model networks (*N*= 1,000) using the function ‘*permatfull’* in the ‘*vegan’* R package *(null model 1)*, keeping the connectance constant but allowing different marginal sums.

Third, we used projected networks to detect cases of extreme reciprocal specializations leading to independent modules. The projected network is the unipartite plant-plant network derived from the bipartite plant-fungus network, where plants interact together if they share at least one fungus. We computed the projected degree, that is the degree of plants in the unipartite network.

To confirm that the patterns of specialization at the global scale hold at a more local scale, we reproduced the analyses of specialization (*k* and *Psp*) in two continental networks in South America and Africa since they represent a high number of interactions and of MH species (Supporting Information Fig. **S5**).

### Phylogenetic distribution of cheating

In order to assess phylogenetic constraints, we explored the phylogenetic distribution of MH cheaters and MH-associated fungi. First, we investigated the phylogenetic distribution of mycoheterotrophy, i.e. if MH plants and MH-associated fungi are more or less phylogenetically related than expected by chance (i.e. patterns of clustering *versus* overdispersion). We computed the net relatedness index (NRI) and the nearest taxon index (NTI) using the ‘*PICANTE’* R package (Kembel *et al.*, 2010). NRI quantifies the phylogenetic structure of a species set based on the mean pairwise distances, whereas NTI quantifies the terminal structure of the species set by computing the mean phylogenetic distance to the nearest taxon of every species (Gotelli & Rohde, 2002). To standardize the indices, we generated 999 null models for each of the following option ‘*taxa.labels’* (shuffles the taxa labels). Significant positive (resp. negative) NRI and NTI values indicate phylogenetic clustering (resp. overdispersion). We computed these indices *(i)* on the plant phylogeny to evaluate the phylogenetic structure of EMH and IMH plants distribution, and *(ii)* on the fungal phylogeny to investigate if MH-associated fungi are phylogenetic structured (we successively tested the distribution of MH-, EMH-, and IMH-associated fungi, and then of the fungi associated with each specific MH family). Similarly, for each plant, we computed the partners’ mean phylogenetic pairwise distance (*MPD*) that is the average phylogenetic distance across pairs of fungal partners (Kembel *et al.*, 2010): a low value of *MPD* indicates that the set of partners is constituted of closely related species. The effect of MH cheating on *MPD* values and its significance was evaluated as for *k* and *Psp* values above.

Second, in order to assess whether fungal partners of a given MH family were derived from fungal partners of AT ancestors or secondary acquired from other MH lineages, we compared in an evolutionary framework the sets of fungi associated with plants with different natures of the interaction. To do so, we computed the unweighted Unifrac distance (Lozupone & Knight, 2005) between sets of fungi interacting with each pair of plants in the network. For each one of the seven MH families, we compared the Unifrac distances across *(i)* every pairs of plant species of this family, *(ii)* every pairs made of one plant of this family and one plant of the most closely related AT family (see Table **2** for AT families), *(iii)* every pairs made of one plant of this family and one plant belonging to other MH families, and *(iv)* every pairs made of one plant of this family and one more distant AT plant (i.e. all AT plants except those of the most closely related AT family). This analysis was not performed on MH Petrosaviaceae, represented by only one species and too divergent to define a reliable AT sister clade.

We tested differences between groups of distances using Mann-Whitney U tests. We also performed a principal coordinates analysis (PCoA) from all the UniFrac dissimilarities of sets of fungal partners, and tested the effect of the nature of the interaction on the two principal coordinates, using Kruskal-Wallis tests. Finally, to examine the extent to which the nature of the interaction predicts fungal partners, we used permutational analysis of variance (PerMANOVA, ‘*adonis’* function in R package ‘*vegan’*), with 10,000 permutations.

## Results

We analyzed the global arbuscular mycorrhizal network filtered from the MaarjAM database (490 plants, 351 fungi, and 26,350 interactions) combined with the reconstructed phylogenetic trees of both partners (Fig. **2**). We estimated a total number of 373 +/− 9 fungi (Chao index), which indicates that the 351 fungi in the dataset included most of the arbuscular mycorrhizal fungal diversity (94% +/− 2%; Supporting Information Fig. S1).

**Figure 2:**
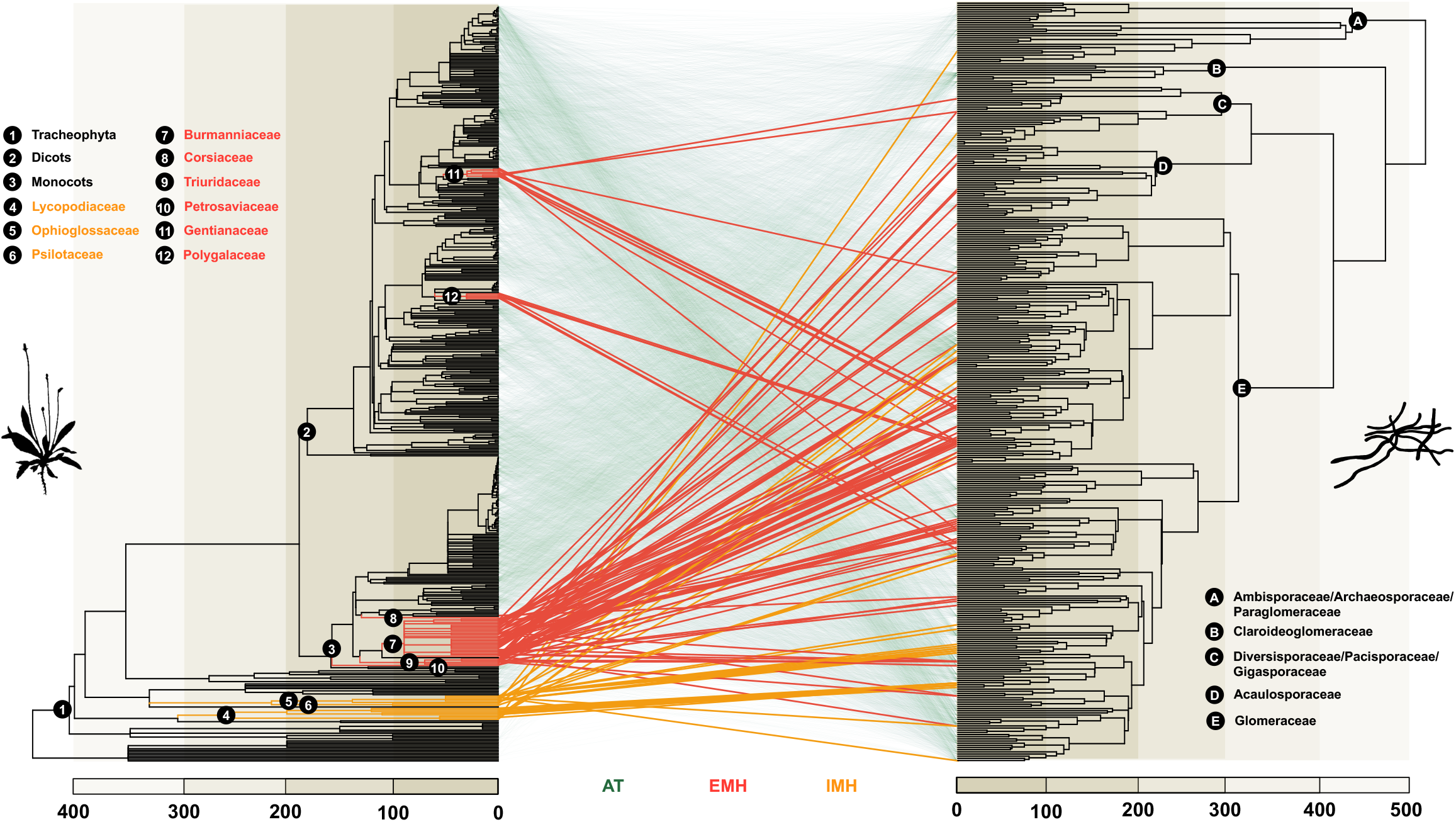
Phylogenetic distribution of mycoheterotroph (MH) cheating in the global arbuscular mycorrhizal mutualism. Phylogenetic trees of 390 plants (left side) and 351 fungi (right side) forming 26,350 interactions (links) in MaarjAM database. Links are colored according to the autotrophic (green), entirely mycoheterotrophic (red), or initially mycoheterotrophic (orange) nature of the plant. Major plant and fungal clades are named. MH cheaters encompass 41 EMH spp. in 6 families, and 15 IMH spp. in 3 families. Scales of the phylogenetic trees are in million years (Myr).

### Network nestedness and specialization of cheaters

The overall network had a significant positive nestedness value (Z-score=9.2, P=1.10^−20^, Supporting Information Table **S4**). Nestedness increased when considering only AT plants (Z-score=16.6, P=8.10^−62^), whereas it was not significant in the network of only MH plants (Z-score=1.44, P=0.075): MH plants reduced nestedness, signifying that they display higher reciprocal specializations. In addition to a main module encompassing most species (481 out of 490 plants and 346 out of 351 fungi), we found three small independent modules (Supporting Information Table S3): *(i)* 6 IMH Lycopodiaceae plants and 3 exclusive fungi (*Glomus* VT127, VT158, VT394); *(ii)* two AT plants from salt marshes (*Salicornia europaea* and *Limonium vulgare*) with one *Glomus* (VT296); and *(iii)* the EMH *Kupea martinetugei* with a unique *Glomus* (VT204).

From the degrees (*k*), we found that EMH and IMH plants are significantly more specialized than AT plants by interacting with on average more than five time less fungi (P=1.4.10^−17^, Kruskal-Wallis test; Fig. **3a**, Table **1**). Partner specializations (*Psp*) indicated that MH plants interact with more specialized fungi (MH-associated fungi themselves interact on average with two times fewer plants; P=5.6.10^−11^; Fig. **3a**). We found similar evidence for MH reciprocal specializations by reanalyzing the network excluding the family Lycopodiaceae (Table **1**; significance assessments using null models are in Supporting Information Table **S5**). This pattern of reciprocal specialization of MH plants and their associated fungi held at a smaller geographical scale in the African and South American networks (Supporting Information Fig. **S5**; Supporting Information Table **S8**; yet the difference was not significant for *Psp* in the South American network, probably due to the small number of species and the low power of the statistical tests).

**Figure 3:**
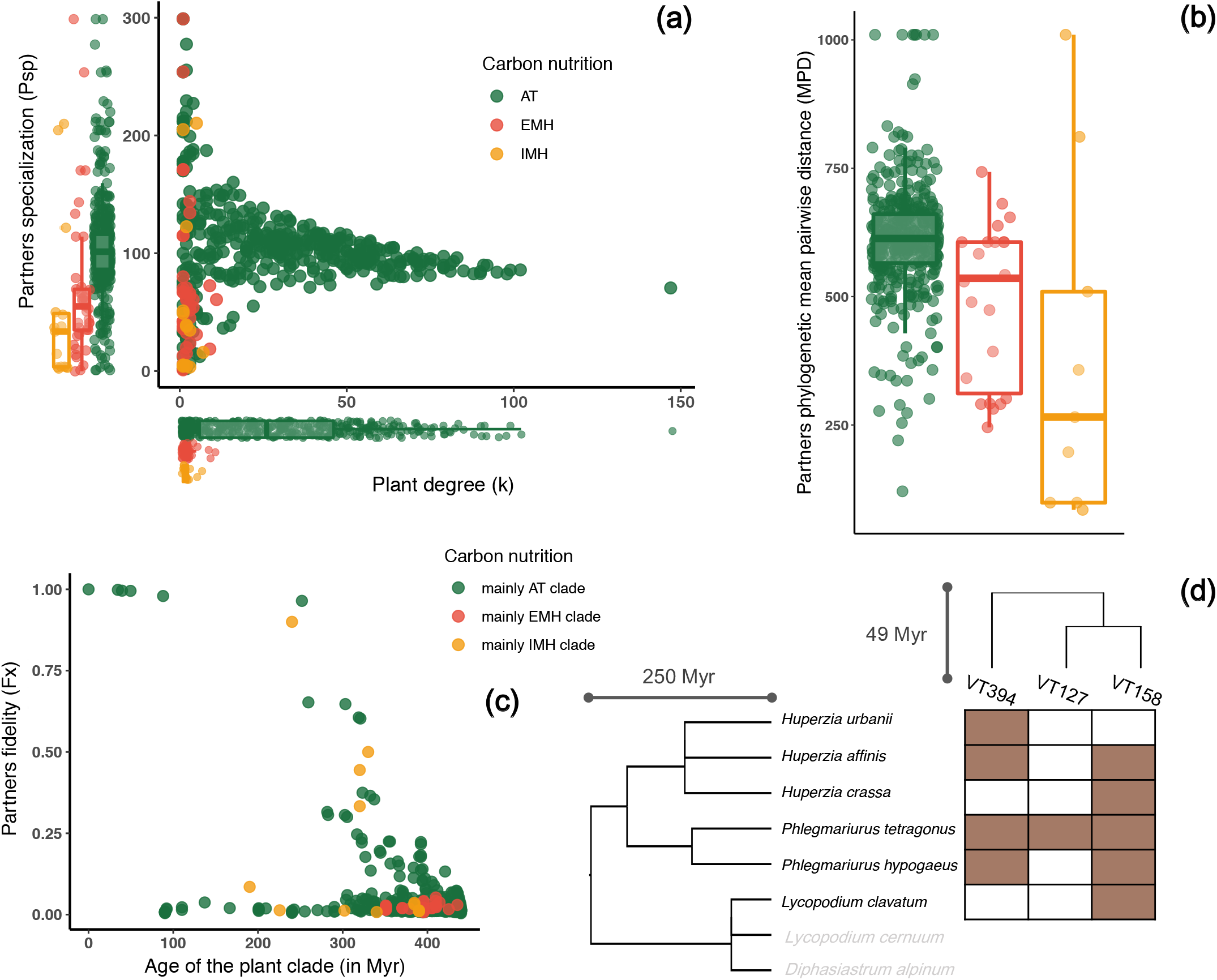
Effect of the nature of the interaction on specialization (*k* and *Psp*), partner’s mean phylogenetic distance (*MPD*), and partner fidelity (*Fx* - see *List of abbreviations* in Supplementary material): (Categories are defined according to the plant carbon nutrition modes, i.e. AT: autotrophic; EMH: entirely mycoheterotrophic over development; and IMH: initially mycoheterotrophic in the development). **(a):** Plant degree (*k*) against fungal partner specialization (*Psp*) (i.e. the average degree of fungal partners); dots in the bottom left corner indicate reciprocal specialization. **(b):** Mean phylogenetic pairwise distance (*MPD*) of the sets of fungal partners. Boxplots present the median surrounded by the first and third quartile, and whiskers extended to the extreme values but no further than 1.5 of the inter-quartile range. **(c):** Fidelity (*Fx*) toward fungal partners in relation to the age of the plant clade. Clades are defined according to their main carbon nutrition mode of their plants (over 50%). The yellow dots departing from other MH clades (high *Fx* values) correspond to clades of Lycopodiaceae. **(d):** Independent network between of the clubmoss family Lycopodiaceae (rows) and their three arbuscular mycorrhizal fungi (columns) with their respective phylogenetic relationships.

**Table 1:** Effect of the nature of the interaction (i.e. plant carbon nutrition modes) on indices of network structure and phylogenetic distributions. The second column corresponds to *P*-values of Kruskal-Wallis tests for the overall network with, or without (in brackets), the Lycopodiaceae. The three last columns correspond to *P*-values of Whitney U tests (pairwise tests) for the overall network including the Lycopodiaceae. *P*-values less than 5% (significance level) are bolded.

| Index | Kruskal-Wallis test | Whitney U tests |  |  |
| --- | --- | --- | --- | --- |
|  |  | AT vs. EMH | AT vs. IMH | IMH vs. EMH |
| Plant degree (*k*) | **1.1e-19 (1.4e-17)** | **5.3e-16** | **4.2e-7** | 0.97 |
| Fungal partner specialization (*Psp*) | **5.6e-11 (1.3e-8)** | **1.1e-4** | **4.5e-9** | 0.054 |
| Mean phylogenetic pairwise distance of fungal partners (*MPD*) | **1.2e-4 (2.0e-3)** | **8.0e-4** | **6.8e-3** | 0.11 |

The partner fidelity index (*Fx*) showed that very few plant and fungi clades interact with ‘clade-specific’ partners (i.e. *Fx*>0.5), and most fungi are shared between different plant clades (Fig. **3c**). Among exceptions, however, the clade of IMH Lycopodiaceae was characterized by a high partner fidelity index (*Fx*>0.8), reflecting a strong association with a clade of 3 Lycopodiaceae-associated fungi (Supporting Information Fig. **S4**). Thus, not only these 6 Lycopodiaceae species and their fungal partners form an independent module, but also the Lycopodiaceae-associated fungi form a monophyletic clade within Glomeromycotina. The estimated clade age was 250 Myr for the Lycopodiaceae and 49 Myr for the Lycopodiaceae-associated fungi (Fig. **3d**), which diverged 78 Myr ago from the other *Glomus* fungi.

### Phylogenetic distribution of cheating

The partners’ mean phylogenetic pairwise distance (*MPD*) indicated that both EMH- and IMH-associated fungi (or even all MH-associated fungi) are phylogenetically more closely related than AT-associated fungi (P=1.6.10^−6^; Table **1**; Fig. **3b**). NRI and NTI values (Supporting Information Table **S7**) also confirmed a significant clustering of MH-, EMH- and IMH-associated fungi on the fungal phylogeny; this clustering held at the family level for fungi associated with each of four main MH families (namely Burmanniaceae, Triuridaceae, Polygalaceae, and Ophioglossaceae). On the plants side, only the EMH plants were significantly clustered, mainly because they all are angiosperms and mostly monocots, but this does not apply for MH plants in general, nor for IMH plants (Supporting Information Table **S7**). These phylogenetic clusters are visually noticeable on fungal and plant phylogenetic trees (Supporting Information Fig. **S2** and **S3**). This demonstrates that although MH cheating evolved several times independently in plants, MH plants interact mainly with closely related fungi (see also Fig. **2**).

Looking specifically at the fungi shared among EMH and IMH plants highlighted differences between EMH and IMH plants (Table **2**). While the IMH Lycopodiaceae family forms an independent module with three specific *Glomus* VTs, another IMH family Ophioglossaceae also had 2 exclusive fungi (*Glomus* VT134 and VT173) among a total of 15 fungi. When comparing the fungi shared between MH families (Table **2**), mainly two closely related families, Burmanniaceae and Triuridaceae, tend to share some fungi with other MH families.

**Table 2:**
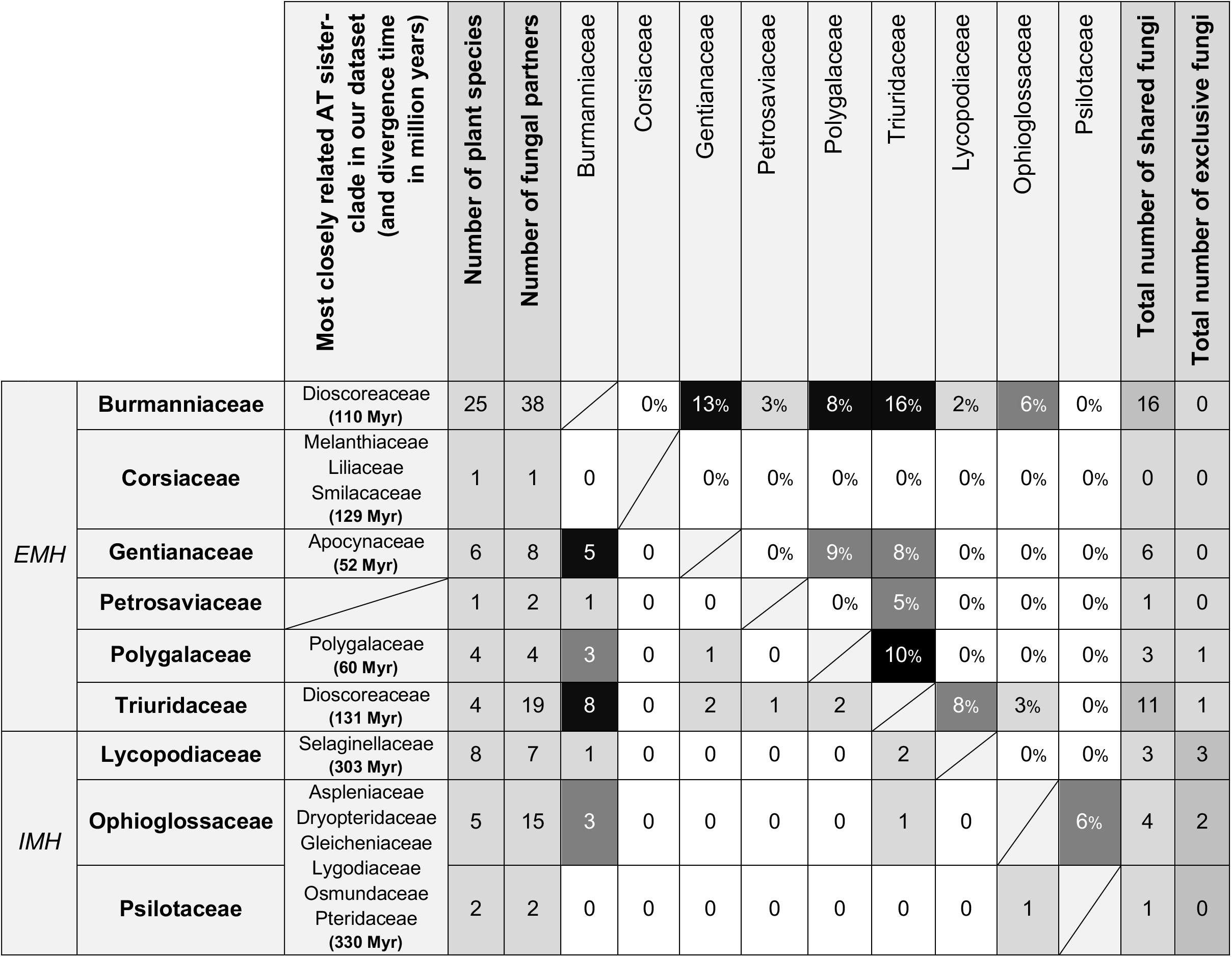
Fungal sharing between nine entirely (EMH) or initially (IMH) mycoheterotrophic plant families. Number (lower part of the matrix) and percentage (upper part) of fungi shared between family pairs. The last two columns represent *(i)* the total number of fungi shared with other EMH or IMH families, and *(ii)* the number of fungi exclusive to this family (i.e. not shared with any other MH or AT family). The second column indicates the most closely related AT sister-clade of each family; it can be either one family, a higher clade, the family itself if AT species were compiled in MaarjAM database (e.g. Polygalaceae), or none in the case of Petrosaviaceae (which forms a too divergent distinct branch).

The decomposition of Unifrac dissimilarities between sets of fungal partners using a principal coordinate analysis, showed a clear pattern of clustering of MH species, supporting that the set of fungal partners associated with MH cheaters are more similar than expected by chance (P<1. 10^−16^ for PCoA1; P= 9.10^−3^ for PCoA2; Fig. **4a**). Similarly, the PerMANOVA analysis indicated that the nature of the interaction (IMH, EMH, or AT) predicts 6.5% of the variance (P=0.0001). By comparing the Unifrac dissimilarities between sets of fungal partners according to the nature of the interaction and plant family relatedness, we observed that all MH families have fungal partners more similar to each other than those of other AT families (Fig. **4b**; Supporting Information Table **S6**). Some families (Burmanniaceae, Polygalaceae, Triuridaceae, Lycopodiaceae, and Ophioglossaceae) had fungal partners significantly more similar to partners interacting with their closest AT relatives (P>0.05) than to partners interacting with other AT families (P<10^−16^). This supports a phylogenetic conservatism of fungal partners during the evolution of MH nutrition in these families. For other MH families (Corsiaceae, Gentianaceae, and Psilotaceae), fungal partners were significantly more similar to partners interacting with other MH families than to partners interacting with their closest AT relatives, the latter being as distant as other AT families (Supporting Information Table **S6**). This supports a shift to new fungal partners correlated with the evolution of MH nutrition in these three families.

**Figure 4:**
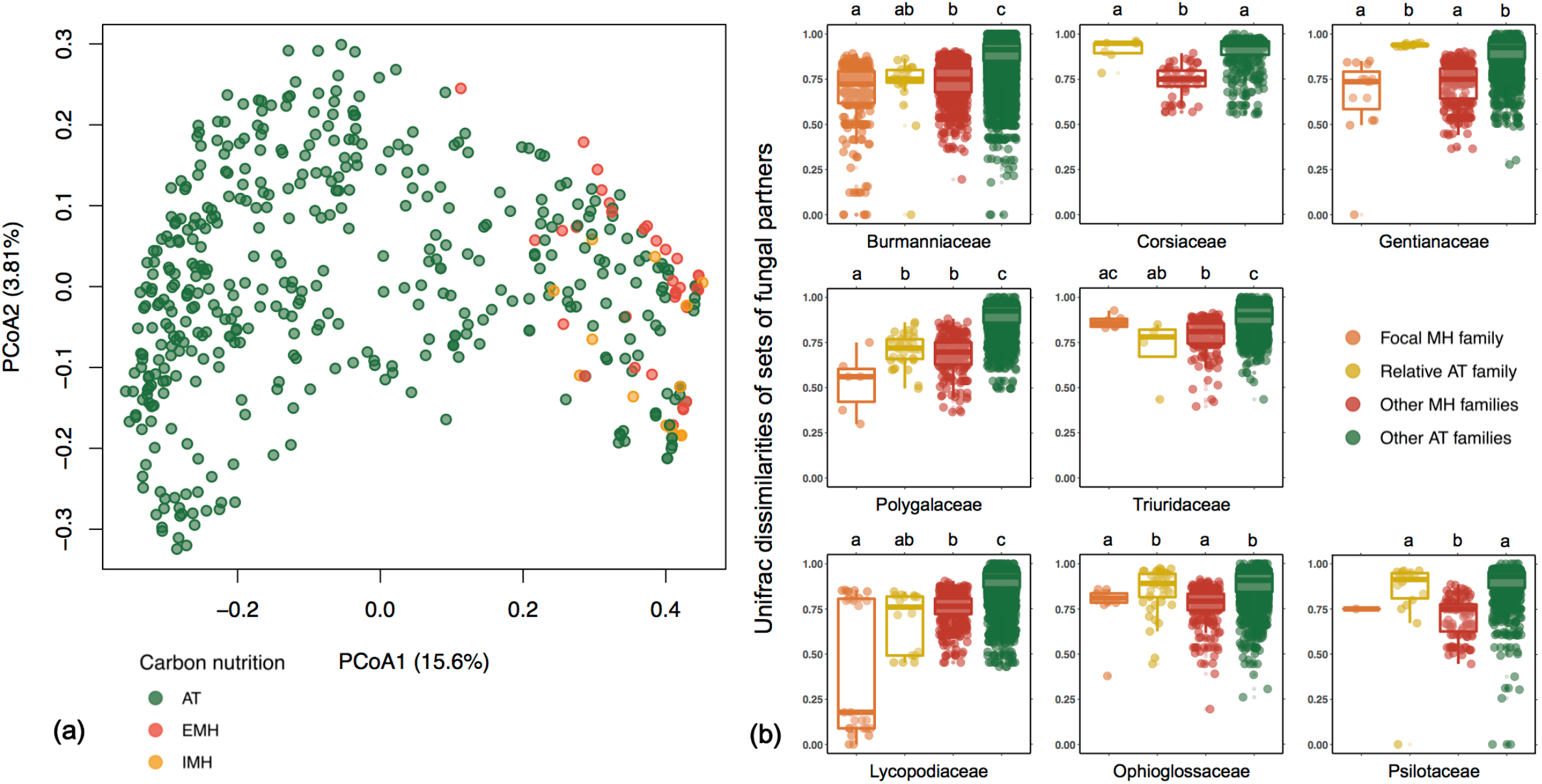
Dissimilarities between sets of fungal partners associated according to the nature of the interaction. **(a):** Principal coordinates analysis (PCoA) from UniFrac dissimilarities of sets of fungal partners. Every dot corresponds to a plant species and is colored according to its autotrophic (green), entirely mycoheterotrophic (red), or initially mycoheterotrophic (orange) nature. Only the first two principal axes explaining respectively 15.6% and 3.8% of the variation were kept. **(b):** Dissimilarities between sets of fungal partners associated with different MH plant families. For each MH family, Unifrac dissimilarities of sets of fungal partners are calculated between one particular MH species belonging to the focal MH plant family and another plant species (from the same family, from the closest related AT family, from other MH families, or from other AT plant families). All the groups cannot be calculated for every MH family, due to the low number of species within families Corsiaceae and Psilotaceae.

## Discussion

We assessed constraints upon the emergence of MH cheating in the mutualistic arbuscular mycorrhizal network. Although this network was nested, we found evidence for reciprocal specialization in the case of MH cheaters (specialists) and their fungal partners (also specialists). We even observed unexpected, extreme reciprocal specialization for some IMH families having exclusive fungi. Finally, we found that independently emerged MH families share many closely related fungi, and that in some of these families fungal partners were derived from AT ancestors, while in others they were acquired by symbiont shift, suggesting different evolutionary pathways leading to MH.

### Cheaters are isolated by reciprocal specialization

We confirmed that MH plants are more specialized toward few mycorrhizal partners than AT plants (Merckx *et al.*, 2012), and showed for the first time that their fungal partners are more specialists than fungi associated with AT plants. This reciprocal specialization is not strict (with the exception of Lycopodiaceae, see below), since MH cheaters and their fungal partners need some connection to AT plants for carbon providing. These findings contrast with the general prediction that, in stable and resilient mutualistic networks, specialists should interact with generalist partners (asymmetrical specialization), resulting in nested networks. We indeed find that MH cheaters lower nestedness in the arbuscular mycorrhizal network. The observed trend toward reciprocal specialization suggests that cheating is an unstable ecological and evolutionary strategy, which could explain the relatively recent origin of MH clades (Fig. **2**). Whatever its origin, the reciprocal specialization of cheaters and their partners was also suggested in other mutualisms (Genini *et al.*, 2010). A parasitic nature of EMH plants has often been guessed (Bidartondo, 2005; Merckx, 2013) but without direct support, in the absence of data on fitness of fungal partners and AT plants linked to MH plants (van der Heijden *et al.*, 2015). Our analysis *a posteriori* supports the view of EMH plants as parasitic cheaters.

There are several non-exclusive explanations to this reciprocal MH specialization. First, functional constraints may act since conditional investment and partner choice occur in the mycorrhizal symbiosis (Kiers *et al.*, 2011): each partner rewards and interacts specifically with the most mutualistic of the many partners they encounter in soil. MH cheaters were able to successfully avoid these constraints with only few specific MH-susceptible fungi with which they now faithfully interact through a parasitic specialization (Selosse & Rousset, 2011). On the fungal side, we can speculate that ‘cheated’ MH-susceptible fungi that provide MH plants with carbon have greater carbon costs for AT plants than other fungi, and that AT plants therefore avoid interactions with these fungi. This would result in reciprocal specialization, and the partial isolation of MH cheaters and their fungal partners from the mutualistic network. The fact that MH plants tend to occur in local patches of low fertility (Gomes *et al.*, 2019) could suggest that in such habitats where access to essential mineral nutrients is limiting, it is even advantageous for AT plants to interact with poorly rewarding fungal partners such as those associated with MH plants.

Second, if it exists a negative trade-off between the cooperativeness of a fungus and its susceptibility to MH plants (for unknown physiological reason, e.g. linked to mechanisms of carbon exchange), cheating MH plants would mainly interact with poorly cooperative fungi, which may be therefore intrinsically avoided by autotrophs.

An in-depth sampling of mycorrhizal interactions and partners’ abundances in various MH local communities would be required to confirm that reciprocal specialization indeed occurs in local communities. Indeed, we observed reciprocal specialization in a large-scale interaction network compiled from mycorrhizal interactions described in different ecosystems around the world, not in locally described physical mycelial networks. This allowed analyzing a global ecological pattern, representing the complete evolutionary history of the partners, and is justified by the very low endemism of arbuscular mycorrhizal fungi and thus the absence of strong geographic structure (Davison *et al.*, 2015; Savary *et al.*, 2018). It is noteworthy that similar patterns of specialization were found in the African and South American networks during this study (Supporting Information Fig. **S5**). On the other hand, a species may appear relatively more specialist in a global network than it actually is in local communities. Reciprocal specialization at the local scale would still need to be confirmed.

### Independent emergences of EMH cheating converge on closely-related susceptible fungi

MH cheating emerged multiple times at different positions of the phylogeny of vascular land plants, indicating weak phylogenetic constraints. This likely results from the low specificity in the arbuscular mycorrhizal symbiosis allowing convergent interactions (Bittleston *et al.*, 2016) in different plant clades. This would have happened during the evolution of MH cheaters with similar MH-susceptible fungi. Thus, functional constraints appear as the main barrier to cheating emergences in the arbuscular mycorrhizal mutualism.

There are however phylogenetic constraints on the fungal side. We found only few fungal clades interacting with independent MH lineages, and they were phylogenetically related in this study as in (Merckx *et al.*, 2012). The susceptibility of fungi to MH cheaters is difficult to discuss, because we know little about physiological and symbiotic traits of arbuscular mycorrhizal fungi (Chagnon et al. 2013; van der Heijden & Scheublin 2007). Yet, interactions between MH cheaters and MH-susceptible fungi likely represent exceptions to the partners choice, which is generally considered to constitutively maintain the stability of this mutualism (Selosse & Rousset, 2011).

The acquisition of MH-susceptible fungi depends on the MH family. In MH families such as Burmanniaceae, fungal partners are closely-related to the fungal partners of AT relatives, suggesting that the MH-associated fungi are derived from the fungal partners of cooperative AT ancestors. In other MH families, such as Gentianaceae or Corsiaceae, fungal partners are more closely-related to fungal partners of other MH families than to AT relatives, suggesting that the MH-associated fungi were secondarily acquired rather than derived from the partners of AT ancestors. A few MH lineages lack closest AT relatives in our analysis (e.g. MH Gentianaceae should be compared to AT Gentianaceae, which are not represented in the MaarjAM database), which may bias our analyses towards supporting secondary transfer from other MH plants rather than acquisition from AT ancestors. Still, similar fungi were found in MH Burmanniaceae and their closest AT relative after a 110 Myrs-old divergence, while MH Gentianaceae and their closest AT relative have distinct fungal partners after a divergence of only 52 Myrs.

Interestingly, all EMH families are evolutionarily relatively recent: the oldest monocot EMH families, such as Burmanniaceae and Triuridaceae, are only 110-130 Myr old, and the dicot EMH families Gentianaceae and Polygalaceae are even more recent (around 50-60 Myr; Zanne *et al.*, 2013; Fig. **2**; Table **2**). The oldest MH families show conservatism for fungal partners while the most recently evolved ones display secondary acquisition. We can speculate that mycoheterotrophy initially emerged in the monocot clade thanks to suitable cheating-susceptible fungal partners; more recently evolved EMH plant families (especially the dicots) then convergently reutilized these fungal partners. Complementary analyses including more sampling of the MH families and their closest AT relatives would be needed to test this speculation.

### Independent networks and parental nurture in IMH

Our results serendipitously revealed that two IMH families, Ophioglossaceae and Lycopodiaceae, have exclusive mycorrhizal associations: they interact with fungi that do not interact with any other plant family. In these families, the fungi are present during both MH underground spore germination and in roots of adult AT individuals (Winther & Friedman, 2007, 2008). AT adults likely act as the carbon source (Field *et al.*, 2015), part of which is dedicated to the offspring. This further supports Leake *et al.* (2008)’s hypothesis of parental nurture where germinating spores would be indirectly nourished by surrounding conspecific sporophytes. Parental nurture is not universal to all IMH families though; in the IMH Psilotaceae for example, fungal partners are shared with surrounding AT plants (Winther & Friedman, 2009). In IMH independent networks, the overall outcome for the fungus over the plant lifespan may actually be positive: fungi invest in MH germinations that represent future carbon sources (Field *et al.*, 2015). In other words, IMH plants do not cheat their exclusive fungi, but postpone reward. We note however that the existence of independent networks for these families should be confirmed in studies of local communities.

We found an extreme reciprocal specialization between Lycopodiaceae and a single *Glomus* clade. Contrary to other early-diverging plant clades that tend to interact with early-diverging fungal clades, the Lycopodiaceae (250-Myr-old) associate with a 49-My-old clade that diverged 78 Myr ago from all other *Glomus* (Rimington *et al.*, 2018). Thus, this highly specific interaction results from a secondary acquisition: some species of Lycopodiaceae may have initially developed IMH with a wider set of fungi, and later evolved into a specific mutualistic parental nurture with their exclusive fungi, raising the possibility of a co-evolution between both clades.

## Conclusions

Our analysis of MH cheating in the arbuscular mycorrhizal symbiosis highlights the role of functional constraints in preventing the emergence of cheaters in mutualisms. Partner choices or sanctions lead to reciprocal specialization which breaks the nested network into modules and isolates MH cheaters and their ‘cheating-susceptible’ partners. Phylogenetic constraints occur on the fungal but not the plant side, with independently emerged MH cheaters that convergently interact with closely related fungi. In addition, our results challenge the general cheater status of mycoheterotrophy, highlighting a dichotomy between true MH cheaters and possibly cooperative, IMH systems with parental nurture. Beyond the mycorrhizal symbiosis, we invite to use our combination of network and phylogenetic approaches to evaluate the strength of constraints upon cheating in other multiple-partner mutualisms (e.g. pollination or seed dispersal).

## Acknowledgments

The authors thank members of the INEVEF team at ISYEB and BIODIV team at IBENS for helpful comments on the article. They also thank C. Fontaine for helpful discussions on network analyses.

This work was supported by a doctoral fellowship from the École Normale Supérieure de Paris attributed to BPL and the École Doctorale FIRE - Programme Bettencourt. MÖ was supported by the European Regional Development Fund (Centre of Excellence EcolChange) and the Estonian Research Council (IUT20-28). HM acknowledges support from the European Research council (grant CoG-PANDA).

## Author contributions

All authors designed the study. MÖ gathered the data, BPL performed the analyses and wrote the first draft of the manuscript, and all authors contributed substantially to the writing and revisions.

## Data Availability Statement

All the data used in this work are available in the MaarjAM database (https://maarjam.botany.ut.ee; Öpik *et al.*, 2010).

## Supporting Information

### Supplementary methods

- Details on the phylogenetic reconstructions
- List of abbreviations

**Supplementary table 1:** Data selection from MaarjAM: description of the used filters.

**Supplementary table 2:** Number and percentage of MH plants and their associated fungi.

**Supplementary table 3:** Independent sub-networks (modules) in the overall arbuscular mycorrhizal network and their respective numbers of partners.

**Supplementary table 4:** Effect of MH cheating on the network nestedness.

**Supplementary Table 5:** Null-model to assess the significance of the effect of the nature of the interaction on indices of network structure and phylogenetic distributions.

**Supplementary Table 6:** Pairwise comparisons of Unifrac dissimilarities between set of fungal partners associated with different plant families.

**Supplementary Table 7:** Measure of the phylogenetic distributions of MH cheating: measure of NRI (net relatedness index) and NTI (nearest taxon index).

**Supplementary Table 8:** Effect of the nature of the interaction (i.e. plant carbon nutrition modes) on indices of the network structure in South America and Africa.

**Supplementary Figure 1:** Rarefaction curves representing the number of fungal taxa as a function of the sampling fraction.

**Supplementary Figure 2:** Calibrated phylogenetic tree of the 490 plant species.

**Supplementary Figure 3:** Calibrated phylogenetic tree of the 351 arbuscular mycorrhizal fungi.

**Supplementary Figure 4:** Fidelity of plant partners (*Fx*) according to the age of the fungal clade.

**Supplementary Figure 5:** Effect of the nature of the interaction on specializations in the South American network and the African network.

**Supplementary figure 6:** The global geographic distribution of sampling sites used in our analysis.

